# Biomechanical control of lysosomal secretion via the VAMP7 hub: a tug-of-war mechanism between VARP and LRRK1

**DOI:** 10.1101/159285

**Authors:** Guan Wang, Sébastien Nola, Simone Bovio, Maïté Coppey-Moisan, Frank Lafont, Thierry Galli

## Abstract

The rigidity of the cell environment can vary tremendously between tissues and in pathological conditions. How this property may affect intracellular membrane dynamics is still largely unknown. Here, using atomic force microscopy, we found that cells deficient in the secretory lysosome v-SNARE VAMP7 were impaired in adapting to substrate rigidity. Conversely VAMP7-mediated secretion was stimulated by more rigid substrate and this regulation depended on the Longin domain of VAMP7. We further found that the Longin domain bound the kinase and retrograde trafficking adaptor LRRK1 and LRRK1 negatively regulated VAMP7-mediated exocytosis. Conversely, VARP, a VAMP7- and kinesin 1-interacting protein, further controlled the availability for secretion of peripheral VAMP7 vesicles and response of cells to mechanical constraints. We propose a mechanism whereby biomechanical constraints regulate VAMP7- dependent lysosomal secretion via LRRK1 and VARP tug-of-war control of the peripheral readily-releasable pool of secretory lysosomes.

## Introduction

From the softest tissue like brain (<1kPa) to the hardest like bones (∼100kPa), the elastic modulus of cell environment can greatly vary in the body of mammals. Matrix elasticity was shown to impact the differentiation of stem cells (Engler et al., 2006), cell spreading and morphology and the capacity to migrate (Tzvetkova-Chevolleau et al., 2008). Previous work showed that exocytosis and endocytosis are regulated by cell spreading and osmotic pressure (Gauthier et al., 2011) and membrane tension regulates secretory vesicle docking through a mechanism involving Munc18-a (Papadopulos et al., 2015). The chemistry of the extracellular matrix also greatly varies in the different tissues and plays a role in regulating cell fate, morphology, and migration (Hakkinen et al., 2011). How substrate rigidity sensing may regulate exocytosis, which in turn regulates membrane tension, is still largely unknown. Secretory mechanisms involve SNAREs, the master actors of intracellular membrane fusion (Südhof and Rothman, 2009). Exocytosis involves the formation of a SNARE complex comprising a vesicular SNARE (v-SNARE) on the vesicle side. The clostridial neurotoxin-insensitive VAMP7 mediates lysosomal secretion (Proux-Gillardeaux et al., 2005). Interestingly enough, VAMP7 was shown to play an essential role in cell migration and invasion (Proux-Gillardeaux et al., 2007; Steffen et al., 2008; Williams and Coppolino, 2011). VAMP7 also contributes into the regulation of membrane composition of sphingolipids and GPI-anchored protein (Molino et al., 2015), which in turn modulates integrin dynamic and adhesion (Eich et al., 2016; van Zanten et al., 2009).

Here we took advantage of atomic force microscopy, micropatterned surfaces, pHluorin live imaging of single vesicle exocytosis (Balaji and Ryan, 2007) and substrate of controlled rigidity and composition to explore the role of lysosomal exocytosis in cell response to biomechanical constraints. Our results suggest that VAMP7-dependent lysosomal secretion responds to rigidity via control by its partners LRRK1 and VARP of the peripheral readily-releasable pool of lysosomes.

## Results

### VAMP7 is required for fibroblast mechano-adaptation

In order to understand the potential regulation of VAMP7 by the cell environment and its potential role in the cell response to mechanical constraints, we first localized VAMP7 in COS7 cells grown on micropatterned glass coverslips. We used cells grown on O pattern, a pattern with homogenous mechanical constraint as control, and Y pattern, a condition where cells are under peripheral traction forces (Albert and Schwarz, 2014). We found that VAMP7 was particularly enriched in actin-rich cell protrusions (Figure 1A and 1B), where contractile forces are generated, in cells grown on a Y pattern. We generated a VAMP7 knockout (KO) COS7 cell line using CRISPR/Cas9 approach (Figure S1A) and measured the cell elasticity on of control and VAMP7 KO COS7 cells by atomic force microscopy (Figure 1C). The cells were grown on soft gels with a rigidity of 1.5 and 28 kPa, two conditions which induce cells to adapt their internal stiffness to the substrate rigidity (Solon et al., 2007). The VAMP7 KO cells reacted differently than control cells to the difference in substrate rigidity, with KO cells presenting a lower elastic modulus when plated on a more rigid substratum at opposite with the control cells (Figure 1D). VAMP7 KO cells also appeared harder despite the environment. This suggested that VAMP7 is required for the proper cell response to environment changes in rigidity. The level of expression of VAMP7 further appeared to have a complex effect on the peripheral positioning of CD63 a marker of secretory lysosomes. Indeed, while KO of VAMP7 did not significantly affect CD63 subcellular localization compared to control cells on Y micropatterns, re-expression of the protein in KO cells modified the distribution of CD63 with an enrichment in cell necks (Figure 1E and 1F). Altogether, these experiments show that VAMP7 is required for proper cell response to biomechanical constraints.

**Figure 1.**
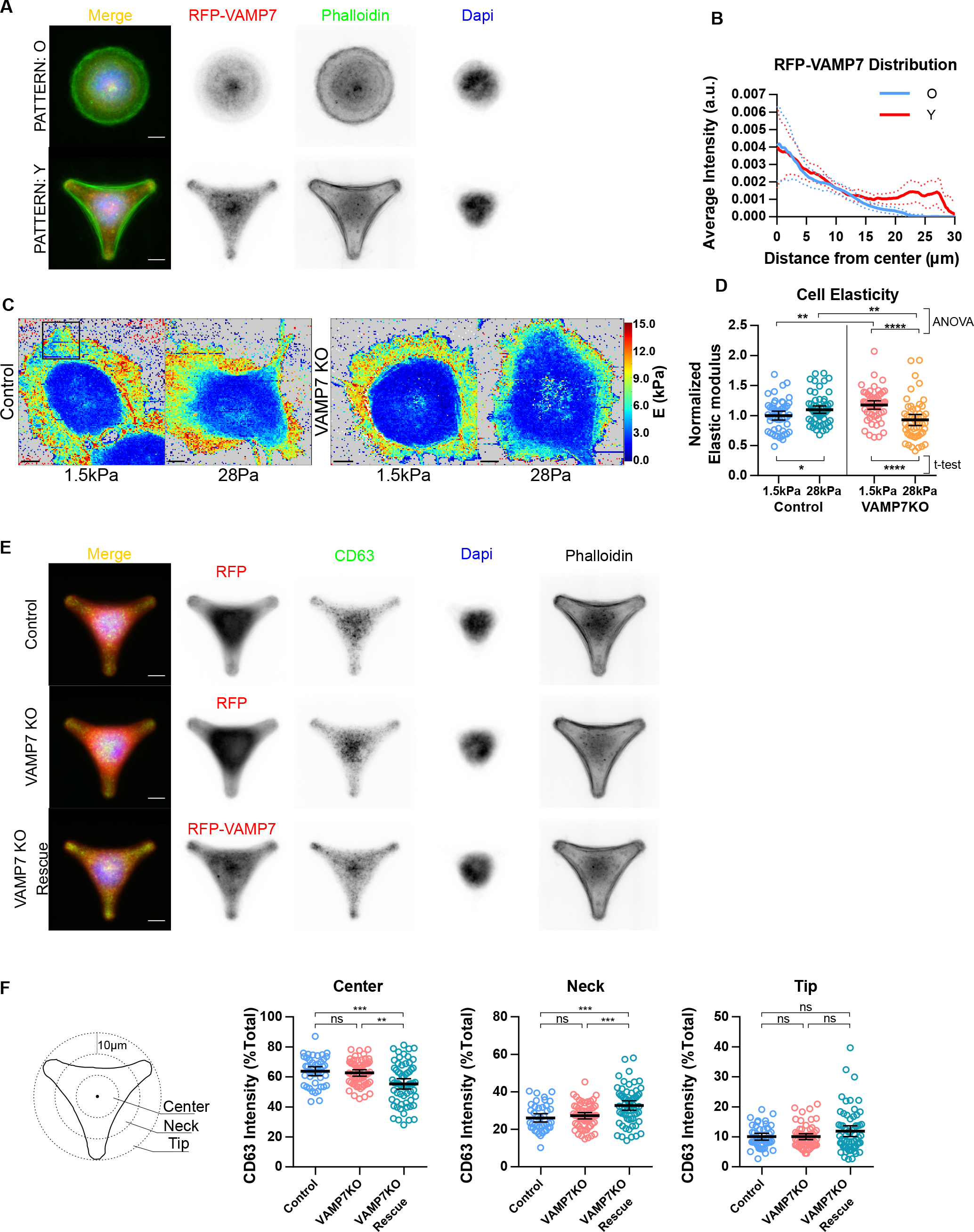
VAMP7 is required for fibroblast mechano-adaptation. (A) Projection of COS7 cells plated on micropattems. n: O=27, Y=27 cells. Scale bar, 10µm. (B) Quantification of RFP-VAMP7 intensity from cell center to cell periphery. Graph shows mean ± 95% CI (dash lines). (C) Heatmaps of cell elasticity plated on laminin coated PDMS gels of 1.5 kPa or 28 kPa. Measurements were systematically made in a 20pm width rectangle area whose typical placement was indicated by the black box. Scale bar, 10µm. (D) Quantification of cell elastic modulus E. Graph shows scatter plot with mean ± 95% CI. Each point represents the median E value of a cell pooled from four independent experiments. *p<0.05, **p<0.01, and ****p<0.0001, ANOVA with Tukey’s post hoc or Welsh’s t-test was used as indicated. (E) Projection of control, VAMP7 KO, and VAMP7 KO re-expressing GFP-VAMP7 cells plated on Y micropatterns. N=43, 59, 63 cells respectively. Scale bar, 10µm. (F) Quantification of CD63 immunofluorescence in cell center area (<10µm from the geometry center), neck area (between 10um and 20µm) and tip area (>20µm). Graph shows scatter plot with mean ± 95% CI. Each point represents the value obtained from cells from two independent experiments. **p<0.01 and ***p<0.001, ANOVA with Tukey’s post hoc.

### Longin-dependent regulation of VAMP7 exocytosis by mechanosensing

We then measured the biophysical properties of individual VAMP7 and VAMP2 exocytic events using pHluorin-tagged molecules (Supplemental Video 1, 2) expressed in COS7 cells grown on surfaces of controlled stiffness generated using PDMS gels of 1.5and 28 kPa. We found that the frequency of exocytosis of VAMP7 had an up to ∼1.5-fold increase on 28kPa in the presence of laminin compared to on 1.5 kPa and to the absence of laminin respectively whereas VAMP2 exocytosis was insensitive to both substrate stiffness and chemistry (Figure 2A and 2B). This finding was confirmed using polyacrylamide gels coated by polylysine or laminin with significant stimulatory effect on VAMP7 exocytosis at 28kPa in the presence of laminin compared to 1.5kPa and the absence of laminin (Figure S1B). Then, we asked whether the regulation of VAMP7 exocytosis could be due to the presence of the Longin domain (LD), a main regulator of VAMP7. Indeed, we found that a mutant of VAMP7 lacking the LD (Δ[1-125]-VAMP7) showed increased exocytosis as previously (Burgo et al., 2013) but its exocytic frequency was not affected by the substrate stiffness and chemistry and was already maximal on soft substrate (Figure 2C). We further analyzed the half-life of pHluorin signals which represents the kinetics of fusion pore opening and spreading followed by endocytosis and re-acidification (Figure S1C). VAMP2 and VAMP7 showed no significant difference in signal persistence depending on stiffness and chemistry. Altogether, these data suggest that VAMP7 exocytosis is modulated by substrate stiffness and composition in a LD-dependent manner. This mode of regulation did not appear to affect the mode of fusion (i.e. transient fusion vs full fusion) thus most likely affects the readily-releasable pool size and/or release probability of VAMP7.

**Figure 2.**
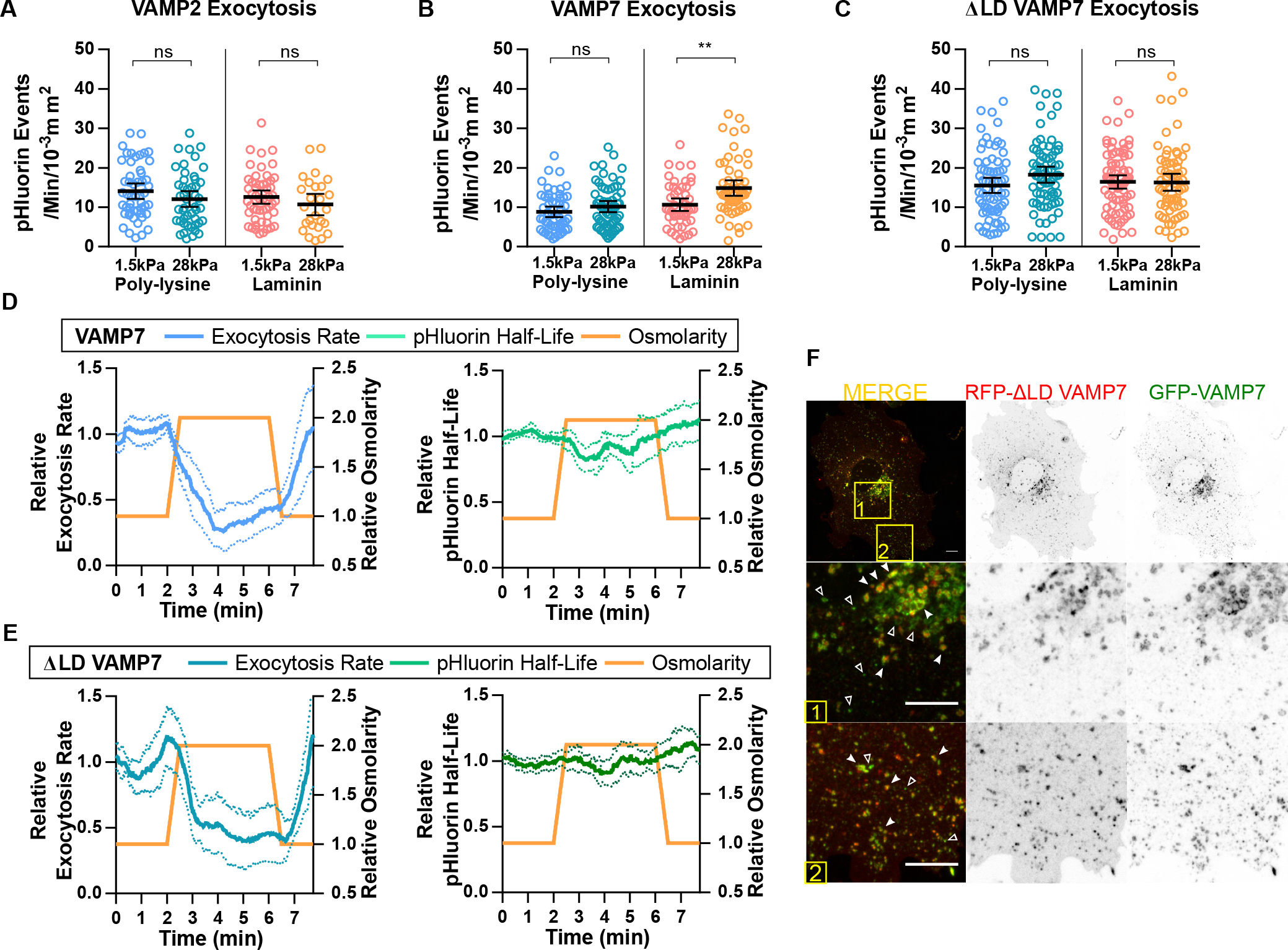
VAMP7-mediated exocytosis is regulated by mechanosensing. (A-C). Quantification of exocytic events in COS7 cells expressing pHluorin-tagged VAMP2, VAMP7 or ΔLD(Δ[1-125]) VAMP7. Cells were plated on poly-lysine or laminin coated PDMS gel of 1.5kPa or28kPa for 18-24 hours. Graph shows scatter plot with mean ± 95%CI. Each point represents the exocytic rate of cells from two or more independent experiments. **p<0.01, Welsh’s t-test. (D and E). Quantification of exocytic rate and pHluorin signals’ half-life in COS7 cells expressing pHluorin-tagged VAMP7 or ΔLD-VAMP7. Cells were plated on laminin coated 28kPa PDMS gels for 18- 24hours. Hyperosmotic shocks were performed by perfusing the 2X osmolality buffer and then washed out by 1X buffer. At each time point, the exocytic rate in the following minute was calculated. Graph shows mean ± 95% CI (dash lines). n>10, pooled from 2 or more independent experiments. (F). Representative COS7 cell co-expressing RFP-tagged ΔLD(Δ[1-120]) VAMP7 and GFP tagged full length VAMP7. Filled arrowheads show the colocalization. Empty arrowheads indicate structures containing only GFP-VAMP7. Scale bar, 10µm.

The previous results suggested that substrate stiffness could have a specific role in VAMP7 regulation. To more directly test the hypothesis of a role of membrane tension in VAMP7 exocytosis, we used hyper-osmotic changes and pHluorin imaging as previously. We found that high hyper-osmotic pressure (2x osmolarity) could instantaneously and reversibly reduce exocytosis frequency of VAMP7 independently of its LD, suggesting different mechanisms of action of membrane tension modulated by osmotic changes and substrate stiffness (Figure 2D and 2E, Supplemental Video 3). We also found that the half-life of pHluorin signals was moderately decreased following hyper-osmotic shocks and then spontaneously restored to normal level. ΔLD-VAMP7 colocalized with full length VAMP7 in the cell periphery but was absent in some perinuclear endosomes (Figure 2F), likely corresponding to late endosomes and lysosomes where VAMP7 is targeted in a LD/AP3-dependent manner (Kent et al., 2012; Martinez-Arca et al., 2003). Therefore, these experiments suggest that substrate rigidity specifically affect lysosomal secretion (VAMP7) and not early endosomal recycling (VAMP2, ΔLD-VAMP7).

Altogether, pHluorin-imaging experiments led us to propose that membrane tension (such as modulated by osmotic shocks) is a master regulator of exocytosis independent of vesicle origin (both endosomal and lysosomal). In the contrary, the regulation of VAMP7 by substrate stiffness appeared not dependent on a pure biomechanical effect via plasma membrane tension but rather required proper sensing of the environment rigidity such as in the presence of laminin.

### The VAMP7 transport hub is regulated by mechanosensing

VAMP7 interactome includes two proteins connected to molecular motors. LRRK1 interacts with VAMP7 through its Ankyrin-repeat and leucine-rich repeat domain and also interacts with dynein (Kedashiro et al., 2015; Toyofuku et al., 2015). VARP interacts with VAMP7 through a small domain in its ankyrin repeat domains and also interacts with kinesin 1 (Burgo et al., 2009, 2012; Schäfer et al., 2012). Interestingly enough, sequence analysis showed that the ankyrin repeat of VARP which interacts with VAMP7 includes a 10aa sequence fully conserved in LRRK1 (Figure 3A). This lead us to wonder whether or not LRRK1 and VARP may participate in the regulation of VAMP7 by substrate stiffness via its LD, in a potentially competitive manner. Firstly, to determine whether or not the interaction between VAMP7 and LRRK1 was through the LD, we carried out in vitro binding assay with GST-tagged cytosolic domain (Cyto), and LD of VAMP7 protein. We found that LRRK1 had a ∼10-fold stronger interaction with LD than with the full-length protein (Figure S2A and S2B). Next, we immunoprecipitated GFP-tagged LRRK1 or GFP-tagged VARP and assayed for coprecipitation of RFP-tagged full length and various deleted forms of VAMP7 from transfected COS7 cells (Figure 3B). We found that LRRK1 interacted with full length, LD and SNARE domain whereas the interaction of VARP was preferentially with full length and SNARE domain, with weak binding to the LD alone (Figure 3C and 3D). The spacer between LD and SNARE domain alone did not bind to either LRRK1 or VARP but appeared to increase the binding of SNARE domain to both LRRK1 and VARP. This likely indicates that the spacer could help the folding of the SNARE domain required for interaction with both LRRK1 and VARP. Nevertheless, the spacer could be replaced by GGGGS motifs of similar length than the original spacer (20aa) without affecting neither LRRK1 nor VARP binding indicating that its role is not sequence-specific but only related to its length. We conclude that LRRK1 interacts with VAMP7 via the LD and its binding to VAMP7 is more sensitive than VARP to the presence of the LD. The loss of mechanosensing of exocytosis when the LD is removed thus likely results from the loss of a competition between LRRK1 and VARP. Unfortunately, with available reagents, competition for binding could not be more directly tested in cells or in vitro. Nevertheless, in good agreement with our hypothesis, triple labelling of exogenously expressed VAMP7, LRRK1 and VARP showed a striking colocalization spots of VAMP7 and VARP in cells tips and colocalization spots of VAMP7 and LRRK1, without VARP, in the cell center (Figure 3F).

**Figure 3.**
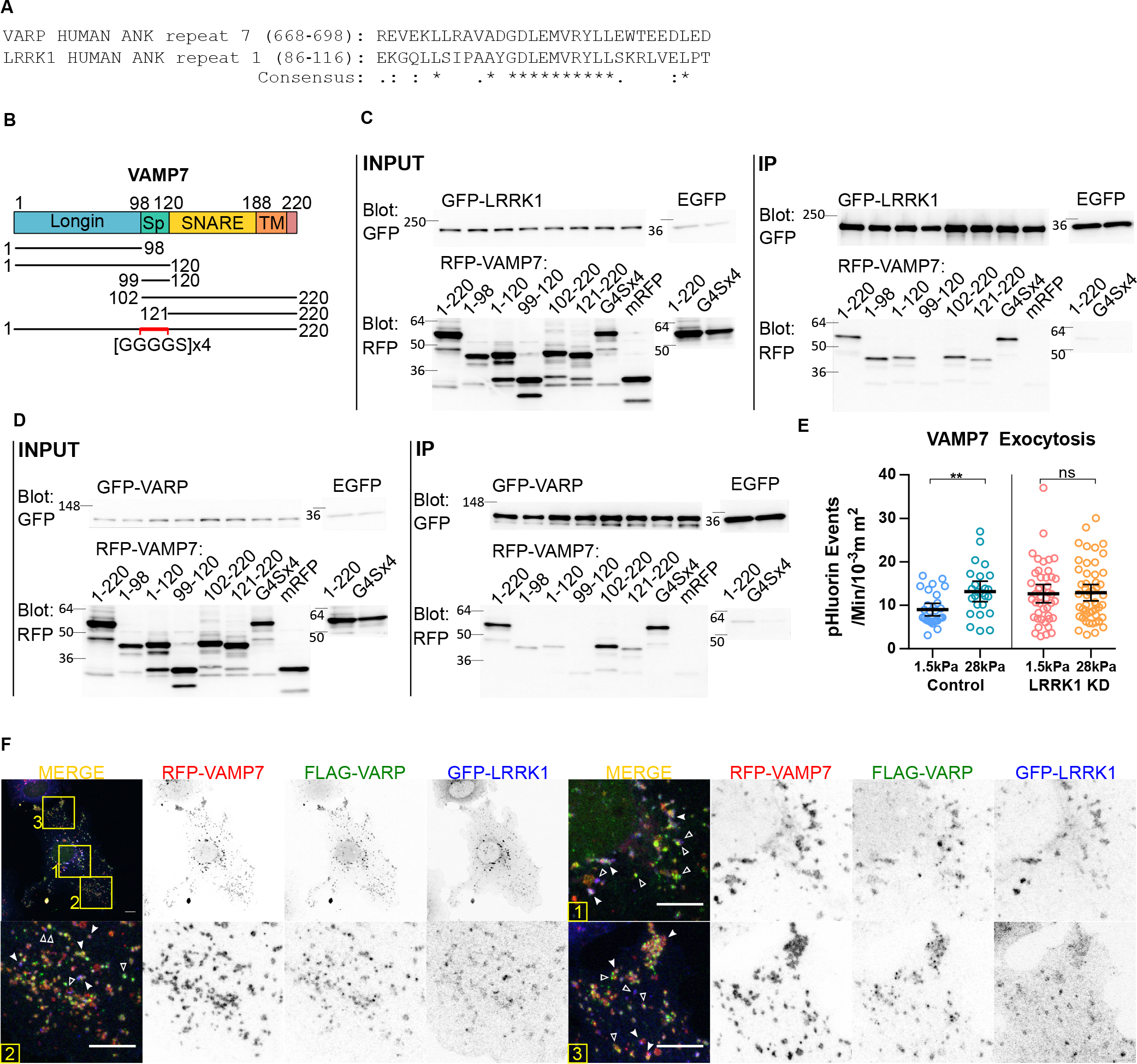
LRRK1 binds VAMP7 and is required for the mechanosensing of VAMP7 exocytosis. (A) Alignment showing that LRRK1 shares a conserved ankyrin repeat domain with VARP, in its interaction domain with VAMP7. (B) Domain organization of rat VAMP7. Sp, spacer; TM, transmembrane. The constructs used for co- immunoprecipitation assay were shown below. (C and D) Assays of binding of LRRK1 and VARP to VAMP7. Lysates from COS7 cells co-expressing GFP-LRRK1 or GFP-VARP with indicated RFP-tagged construction of VAMP7 were immunoprecipitated (IP) with GFP-binding protein (GBP) fixed on sepharose beads. Precipitated proteins were subjected to SDS-PAGE, and the blots were stained with antibodies against indicated target proteins. EGFP protein and mRFP protein were used as control for nonspecific binding. The experiment has been independently repeated three times with similar results. (E) Quantification of exocytic events in COS7 cells co-expressing VAMP7-phluorin with control shRNA or LRRK1-shRNA, growing on laminin coated PDMS gels for 18-24hours. Graph shows scatter plot with mean ± 95% CI. Each point represents the exocytic rate of cells from two independent experiments. **p<0.01, Welsh’s t-test. (F) Representative COS7 cell co-expressing RFP-VAMP7, FLAG-VARP and GFP-LRRK1. Filled arrowheads indicate triple colocalization. Empty arrowheads indicate structures where either FLAG- VARP or GFP-LRRK1 is missing or dominant. Scale bar, 10µm.

To further decipher the role of LRRK1, we silenced its expression by shRNA and assayed for VAMP7 exocytosis on soft and rigid substrate. We found that the exocytosis frequency of VAMP7 on soft substrate was increased to the same level as on rigid substrate in cells in which the expression of LRRK1 was knocked down (Figure 3E). This suggested the striking hypothesis that LRRK1 is indispensable for the sensing of substrate softness.

According to previous work on LRRK1, VAMP7-LRRK1 interaction should recruit CLIP-170 and dynein allowing for retrograde transport on microtubules (Kedashiro et al., 2015). To further understand the potential role of LRRK1 in VAMP7 trafficking, we carried out live imaging of cells expressing GFP- LRRK1 and RFP-VAMP7, and found that VAMP7 and LRRK1 accumulated together in the cell center upon EGF stimulation (Figure S2C and S2D), a condition promotes perinuclear localization of LRRK1- containing endosomes (Hanafusa et al., 2011; Ishikawa et al., 2012). Analysis of confocal images taken from cells expressing GFP-tagged WT LRRK1, Y944F or K1243M mutants (constitutively active and inactive kinase form of LRRK1 respectively) and RFP-tagged VAMP7 showed that VAMP7 accumulated more in the perinuclear region in LRRK1 Y944F expressing cells, and more towards the cell periphery in LRRK1 K1243M expressing cells (Figure S2E and S2F), suggesting that LRRK1 kinase activity enhanced the retrograde transport of VAMP7 vesicles into the perinuclear region. LRRK1 was previously found to play a role in autophagy (Toyofuku et al., 2015) but we did not find significant autophagy induction as seen by LC3-II imaging in cells on soft vs rigid substrates and western blotting (Figure S2G and S2H). We conclude that LRRK1 mediates retrograde transport of VAMP7 in a kinase-dependent activity and that LRRK1 is required for the control of VAMP7 exocytosis in response to substrate rigidity.

### Opposite roles of LRRK1 and VARP in mechanosensing

A prediction from our previous results showing that VAMP7 exocytosis is required for mechanosensing (Figure 1C) and that LRRK1 and VARP generate a tug-of-war mechanism for the cell positioning of secretory lysosomes (Figures 3, S2) would be that LRRK1 and VARP should themselves play a role in mechanosensing. To test this hypothesis, we again used the previous assay with cells grown on substrates of different rigidities. We found that soft substrate promoted more perinuclear accumulation of VAMP7 that rigid substrate (Figures 4A), similar to the effect of LRRK1 Y944F mutant. We found that VAMP7 was localized more to the center in LRRK1-overexpressing cells. The opposite was found in VARP- overexpressing cells which showed decreased center-localized VAMP7. VARP-overexpressing cells further striking concentration of VAMP7 at the tips of cell protrusions. The effects of LRRK1 and VARP overexpression were not sensitive to substrate rigidity. This later data suggests that the effect of overexpression of these proteins dominated over the regulation that occurs between soft and rigid environment when they are expressed at physiological levels. In order to further decipher the role of VARP and LRRK1, we then used Crispr/Cas9 approach to knock out the expression of the proteins (Figure S1A), cultured the KO cells on substrate of 1.5 and 28 kPa, and assayed for perinuclear accumulation of RFP-VAMP7. We again reproduced the decreased perinuclear concentration of VAMP7 on more rigid substrate in control cells. The effect of rigidity was lost in LRRK1 KO cells. In contrary, re-expression of LRRK1 in KO cells exacerbated central concentration of VAMP7 in a rigidity independent manner. Conversely, VARP KO showed a strong perinuclear accumulation of VAMP7 on rigid substrate and this effect was reversed by re-expression of VARP. In this later case, the effect of substrate rigidity was visible after VARP re-expression in VARP KO cells. Altogether, these experiments using KO and overexpression approaches and culture on soft and rigid substrate suggest that LRRK1 and VARP provides a tug-of-war mechanism which mediates the fine tuning of VAMP7 subcellular localization regulated by mechanical constraints. In this regulatory mechanism, the precise expression level of LRRK1 and VARP here appeared to be a critical parameter, further reinforcing the notion of a competitive mechanisms strongly dependent on the concentration and activity of LRRK1 and VARP.

**Figure 4.**
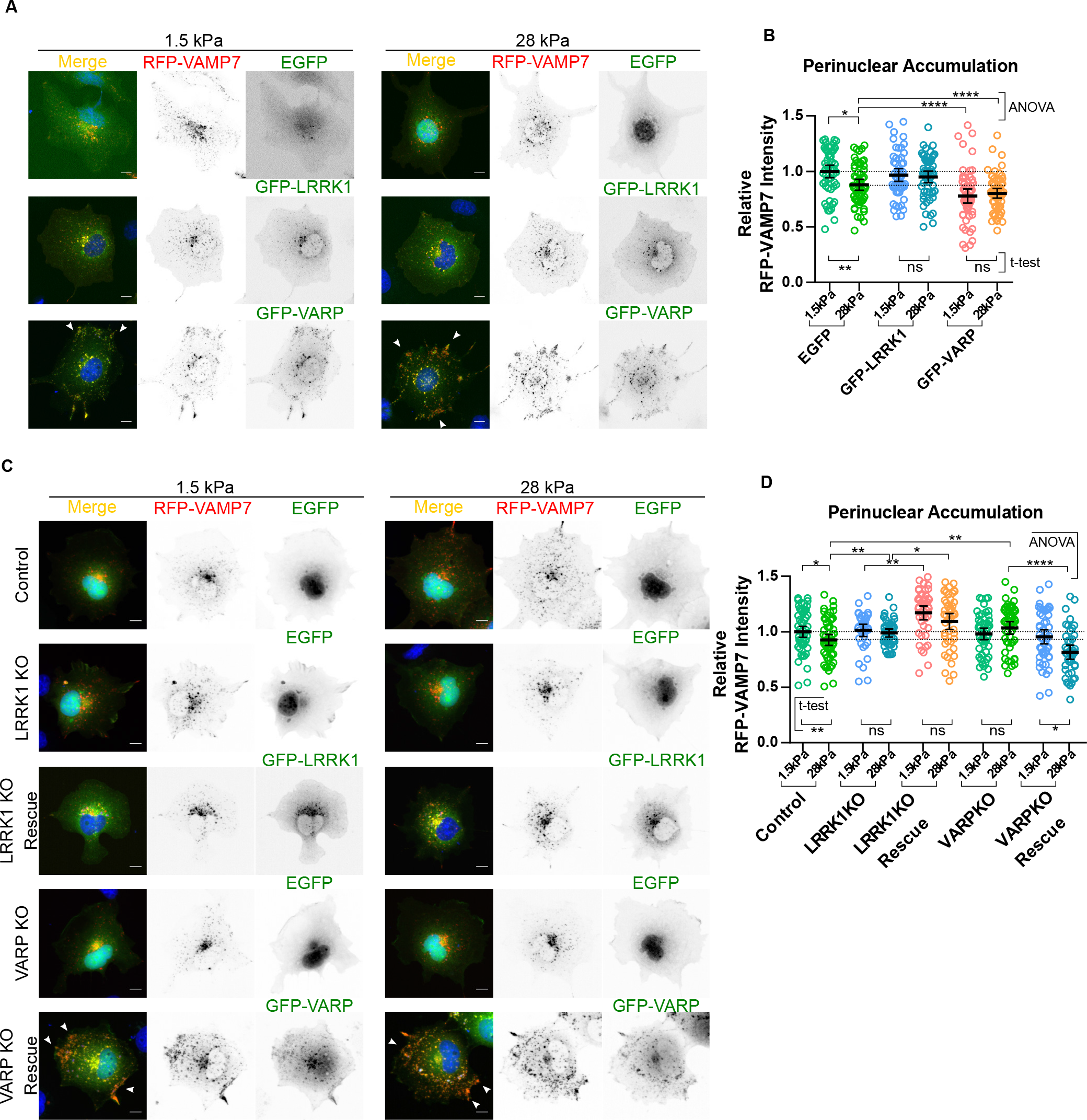
LRRK1 and VARP have opposite roles of in rigidity dependent VAMP7 positioning. (A) Representative WT COS7 cells co-expressing RFP-tagged VAMP7 with GFP-tagged LRRK1 or VARP, growing on laminin coated PDMS gels. (C) Representative Control, LRRK1 KO and VARP KO COS7 cells growing on laminin coated PDMS gels. Control and KO cells were transfected with RFP-VAMP7 and EGFP as indicated, and with GFP- LRRK1 and VARP in rescue conditions. Images show z-projection of confocal stack. Arrowheads show the colocalization in cell protrusions. Scale bar: 10µm. (B and D) Quantification of RFP-VAMP7 fluorescence in the perinuclear region. Graph shows scatter plot with mean ± 95% CI. Each point represents the value obtained from a cell pooled from two independent experiments. *p<0.05, **p<0.01 and ****p<0.0001, ANOVA with Tukey’s post hoc or Welsh’s t-test was used as indicated.

## Discussion

In this study, we found that VAMP7-dependent lysosomal exocytosis was required for cells to sense substrate rigidity and that the latter redistributed VAMP7 to the cell periphery in a LD, VARP- and LRRK1-dependent manner. LRRK1 and VARP, appeared to operate via an opposite control of the availability for secretion of peripheral VAMP7 vesicles in response to mechanical constraints, thus suggesting a tug-of-war mechanism.

VAMP7 KO cells showed increased elastic modulus on soft substrate and a decreased modulus on more rigid substrate compared with control cells. This likely suggest that the lack of VAMP7 may prevent cells from properly responding to mechanical constraints (Figure 1). Conversely, substrate rigidity increased exocytosis of VAMP7, but not VAMP2. This likely indicates the need for different types of membranes being transported to the cell surface depending on the biophysical properties of cell environment, particularly its rigidity. VAMP7 was shown to be important for phagophore formation and autophagosome secretion (Fader et al., 2012; Moreau et al., 2011) and rigidity was shown to increase autophagy (Ulbricht et al., 2013) but we did not find significant LC3-II induction in the different conditions tested so we do not think that substrate stiffness significantly activated autophagy in our experimental conditions. More likely we think our findings are related to the previous demonstration that VAMP7 mediates the transport of GPI-anchored proteins and lipid microdomains to the plasma membrane (Lafont et al., 1999; Molino et al., 2015; Pocard et al., 2007). Accordingly, increased exocytosis of GPI-anchored proteins was found in the secondary contractile phase during cell spreading (Gauthier et al., 2011). An attractive hypothesis would be that VAMP7 bring lipids which best fit a membrane under higher tension such as on more rigid substrates. Further studies are now required to decipher the precise signaling mechanism of how rigidity sensing and cortical tension regulate VAMP7 exocytosis.

We found that substrate rigidity increased and hyperosmotic shock inhibited the exocytosis frequency of VAMP7 following a remarkably quick adaptation of exocytosis frequency to strong changes in membrane tension. The effect of hyperosmotic shock on persistence of the signal at plasma membrane would be best explained by decreased fusion pore flattening because fusion pore growth is promoted or even driven by the membrane tension (Bretou et al., 2014) and potential increased recovery of plasma membrane by endocytosis upon the osmotic shock. Our data thus fit well with the notion that exocytosis increases the surface area therefore decreases membrane tension, thus needs to be shut down to compensate for decreased membrane tension following hyperosmotic shock (Gauthier et al., 2011; Keren, 2011; Sens and Plastino, 2015). Nevertheless, we found similar effects of hyperosmotic shock on VAMP7 deleted of its LD while this was not the case for increased substrate stiffness. This indicates that acute changes of cell tension, such as osmotic shocks, acting likely via a direct effect on membrane tension, and secretory vesicles in close proximity with the plasma membrane, proceed from different mechanisms than substrate stiffness.

The mechanism unraveled here further suggest the involvement of two members of VAMP7’s hub in mechanosensing-dependent regulation of transport and exocytosis. Here we found that LRRK1 strongly interacts with LD and SNARE domain of VAMP7 with a particularly strong interaction with LD *in vitro*. LRRK1 and VAMP7 were co-transported to the cell center upon EGF addition. Silencing LRRK1 removed the regulation of VAMP7 exocytosis by substrate rigidity. LRRK1 overexpression concentrated VAMP7 in the cell center. This effect dominated over substrate rigidity, and was further emphasized by the kinase activity as it was previously shown in the case of the EGFR (Ishikawa et al., 2012). VARP mediates transport of VAMP7 to the cell periphery (Burgo et al., 2009, 2012; Hesketh et al., 2014). Here we found that VARP bound efficiently ΔLD VAMP7 and its overexpression decreased the perinuclear pool of VAMP7 while increasing the peripheral one. Our data thus give a reasonable explanation for the increased exocytosis frequency of ΔLD VAMP7 as the later would still efficiently bind to VARP and less to LRRK1. Altogether, the present data lead us to propose a tug-of-war mechanism with LRRK1 on the retrograde end and VARP on the anterograde end of VAMP7 trafficking. Our results further suggest that substrate stiffness would be able to regulate the tug-of-war between LRRK1 and VARP for lysosome positioning in the cell periphery and exocytosis. The effects of LRRK1 and VARP suggest that their concentration in the cell is important for VAMP7 center to periphery distribution, fitting well with the notion of a tug-of-war mechanism. In conclusion, we suggest that VAMP7 lysosomal secretion is regulated by biomechanical constraints relayed by LRRK1 and VARP, a mechanism with potential broad relevance for plasma membrane dynamics in normal conditions (Koseoglu et al., 2015), infection (Chiaruttini et al., 2016; Larghi et al., 2013) and cancer (Steffen et al., 2008).

## Experimental Procedures

Detailed procedures and reagent information are in the Supplemental Experimental Procedures. GraphPad Prism software were used for statistical analyses. Data were analyzed using Welsh’s t-test or one-way ANOVA followed by a Tukey post hoc test as indicated in legends.

## Author Contributions

Conceptualization, G.W. and T.G.; Methodology, G.W. and S.N.; Software, G.W.; Investigation, G.W., S.N. and S.B.; Writing – Original Draft, G.W. and T.G.; Funding Acquisition, G.W. and T.G.; Supervision, T.G., F.L. and M.C.

## Acknowledgements

We are grateful to Dr. Michael Kozlov (Tel Aviv University, Israel) for critical reading of a preliminary version of the manuscript. We thank V. Proux-Gillardeaux for helpful discussion, C. Vannier and A. Verraes for providing materials. We are grateful to Dr. H. Hanafusa for the LRRK1 vectors. Work in our group was funded by grants from INSERM, CNRS, Association Française contre les Myopathies (Research Grant 16612), the French National Research Agency (Neuro*ImmunoSynapse* ANR-13-BSV2- 0018-02), the Ecole des Neurosciences de Paris (ENP), the Fondation pour la Recherche Médicale (FRM), *Who am I?* Labex (Idex ANR-11-IDEX-0005-01), awards of the Association Robert Debré pour la Recherche Médicale to T.G. and fellowship from the Region Ile-de-France in the framework of DIM Cerveau&Pensée and from FRM (FDT20150532766) to G.W. We acknowledge the ImagoSeine facility, and the France BioImaging infrastructure supported by ANR (10-EQPX-04-01) and the EU-FEDER (12,001,407) as part of “Investments of the future” program.

